# Niclosamide potentiates TMEM16A and induces vasoconstriction

**DOI:** 10.1101/2023.07.31.551400

**Authors:** Pengfei Liang, Yui Chun S. Wan, Kuai Yu, H. Criss Hartzell, Huanghe Yang

**Author notes:** Correspondence (H.Y.).

## Abstract

The TMEM16A calcium-activated chloride channel is a promising therapeutic target for various diseases. Niclosamide, an anthelmintic medication, has been considered as a TMEM16A inhibitor for treating asthma and chronic obstructive pulmonary disease, but was recently found to possess broad-spectrum off-target effects. Here we show that, under physiological conditions, niclosamide acutely potentiates TMEM16A without having any inhibitory effect. Our computational and functional characterizations pinpoint a putative niclosamide binding site on the extracellular side of TMEM16A. Mutations in this site attenuate the potentiation. Moreover, niclosamide potentiates endogenous TMEM16A in vascular smooth muscle cells, triggers intracellular calcium increase, and constricts the murine mesenteric artery. Our findings advise caution when considering niclosamide as a TMEM16A inhibitor to treat diseases such as asthma, COPD, and hypertension. The identification of the putative niclosamide binding site provides insights into the mechanism of TMEM16A pharmacological modulation, shining light on developing specific TMEM16A modulators to treat human diseases.

## INTRODUCTION

TMEM16A, also known as anoctamin-1 (ANO1), is a *bona fide* calcium-activated chloride channel (CaCC) (Caputo et al., 2008; Schroeder et al., 2008; Yang et al., 2008). It is widely expressed in various cell types and plays crucial roles in regulating physiological processes such as smooth muscle contraction, gut motility, fluid secretion, cell volume regulation, nociception, and anxiety (Pedemonte and Galietta, 2014; Whitlock and Hartzell, 2017; Kunzelmann et al., 2019; Hawn et al., 2021; Le et al., 2021; Song et al., 2022). Upregulation of TMEM16A has been reported in a wide range of pathological conditions, including inflammatory airway disease, asthma and chronic obstructive pulmonary disease (COPD), cancer, hypertension, and ischemia/reperfusion injury (Ma et al., 2017; Wang et al., 2017; Crottes and Jan, 2019; Papp et al., 2019; Bai et al., 2021; Korte et al., 2022). Therefore, pharmacological inhibition of TMEM16A provides a promising new strategy to treat these diseases.

A wide variety of TMEM16A inhibitors or antagonists have been identified since its molecular cloning in 2008 (Namkung et al., 2011; Peters et al., 2015; Seo et al., 2016; Centeio et al., 2020; Liu et al., 2021; Al-Hosni et al., 2022). Niclosamide, one of the World Health Organization’s essential medicines for treating tapeworm infections, was recently reported to be a TMEM16A antagonist (Miner et al., 2019; Henckels et al., 2020). This finding subsequently led to the proposal that niclosamide could be repurposed to suppress upregulated TMEM16A activity in inflammatory airway diseases such as asthma and COPD (Cabrita et al., 2019; Ousingsawat et al., 2022). However, other studies showed contradictory results that do not support niclosamide as a clean TMEM16A inhibitor. Niclosamide has off-target effects including disruption of calcium signaling by inhibiting calcium store release (Ousingsawat et al., 2022; Genovese et al., 2023).

To investigate the effects of niclosamide on TMEM16A, here we use patch clamp, molecular docking and simulation, structure-guided mutagenesis, calcium imaging, and pressure myograph to study the effect of niclosamide on heterologously expressed TMEM16A in HEK293T cells and endogenous TMEM16A in vascular smooth muscle cells (SMCs). We find that, under physiological calcium and voltage, niclosamide acutely potentiates TMEM16A activation without any obvious inhibitory effect. Micromolar niclosamide potentiates TMEM16A by binding to a putative extracellular binding site. We further demonstrate that niclosamide potentiates endogenous TMEM16A in vascular SMCs, increases intracellular calcium, and constricts the mesenteric artery. Our study, together with another recent report (Danahay et al., 2023), shows no evidence that niclosamide inhibits TMEM16A CaCC. The acute potentiation effect of niclosamide on TMEM16A imposes a warning about repurposing niclosamide as a TMEM16A inhibitor to treat human diseases such as asthma and COPD. The putative niclosamide binding site identified in this study, on the other hand, may inform the development of more potent and specific TMEM16A potentiators to mitigate diseases that require augmented CaCC activity, including cystic fibrosis and gastroparesis.

## RESULTS

### Niclosamide potentiates exogenously expressed TMEM16A

To elucidate the effects of niclosamide on TMEM16A current, we conducted whole cell patch clamp recording of HEK293T cells overexpressing TMEM16A (Fig. 1). With a physiological intracellular Ca^2+^ concentration of 500 nM and a −60 mV holding voltage, exogenous TMEM16A gave rise to characteristic outwardly rectifying currents with slow activation and deactivation kinetics (Fig. 1B). Upon extracellular application of 5 µM niclosamide, the TMEM16A channel appears to become constitutively open. TMEM16A current was potentiated at both positive and negative voltages (Fig. 1A), but inward current was potentiated to a greater extent than outward current so that the current-voltage relationship became nearly linear, unlike the outward rectification typically observed at this Ca^2+^ concentration. (Fig. 1B, D). These changes resemble the TMEM16A current at saturating levels of Ca^2+^ (Xiao et al., 2011; Huang et al., 2012). These effects were observed in the authors’ two laboratories independently using different TMEM16A splicing isoforms (see Methods for details), ruling out potential systemic error (Fig. 2). Our analysis of the conductance-voltage (*G-V*) relationships normalized to the maximum tail current amplitude at +140 mV before niclosamide application further demontrates the potentiation (Fig. 1C). Because the extracellular solution in this experiment contained 5 mM EGTA to chelate free Ca^2+^, we exclude the possibility that the potentiation is caused by Ca^2+^ influx. An effect of niclosamide on Ca^2+^ release from stores is excluded by the Ca^2+^ buffering of the pipet solution. The potentiation effect of niclosamide is dose-dependent with an EC_50_ (half-maximal effective concentration) of 1.16 ± 0.16 µM (Fig. 1D-E) and is completely reversible (Fig. 1A-C). Our whole-cell patch clamp experiments thus demonstrate that niclosamide potentiates exogenous TMEM16A CaCC under physiological conditions.

**Figure 1.**
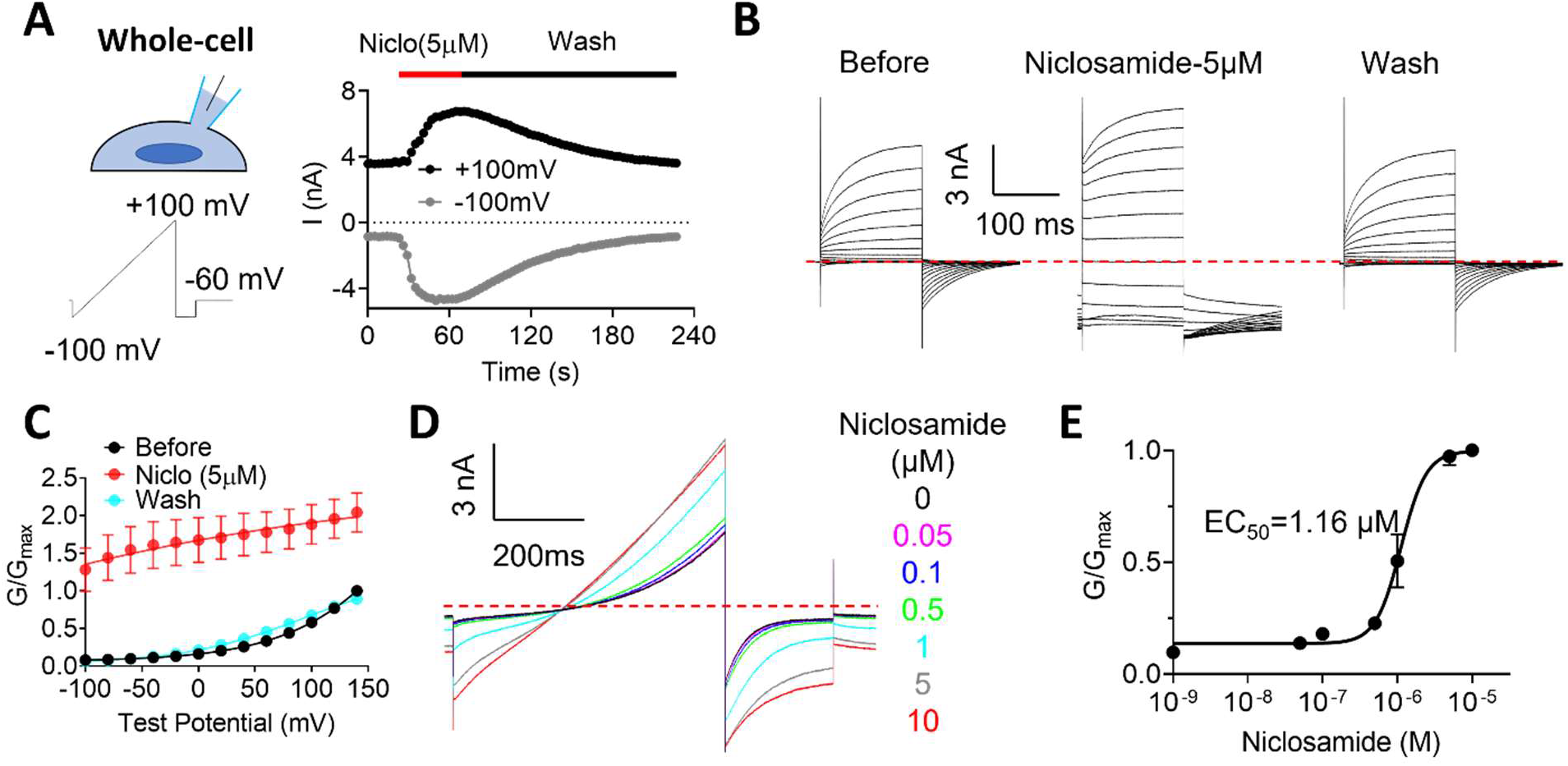
Niclosamide activates TMEM16A exogenously expressed in HEK293T cells. **A.** Representative whole-cell current recorded in HEK293T cells overexpressing TMEM16A ‘a’ splicing isoform. Intracellular solution contains 500 nM free Ca^2+^. The current was elicited by a ramp protocol from −100 mV to +100 mV. Holding potential was −60 mV. The patch clamp configuration and voltage protocol are shown on the left. **B,** Representative current traces elicited by a voltage-step protocol from −100 mV to +140 mV with 20-mV increments. Holding potential was −60 mV. **C,** *G-V* relationship of TMEM16A current recorded in B. Error bars represent SEM, n=7. The tail currents were normalized to the maximum tail current at +140 mV before niclosamide. **D,** Representative current traces in response to different niclosamide concentrations as indicated. The current was elicited by a ramp protocol the same as in panel A. **E,** Niclosamide dose-response curve. The data were fitted to the Boltzmann equation with EC_50_ of 1.16 µM. Error bars represent SEM, n=5.

**Figure 2.**
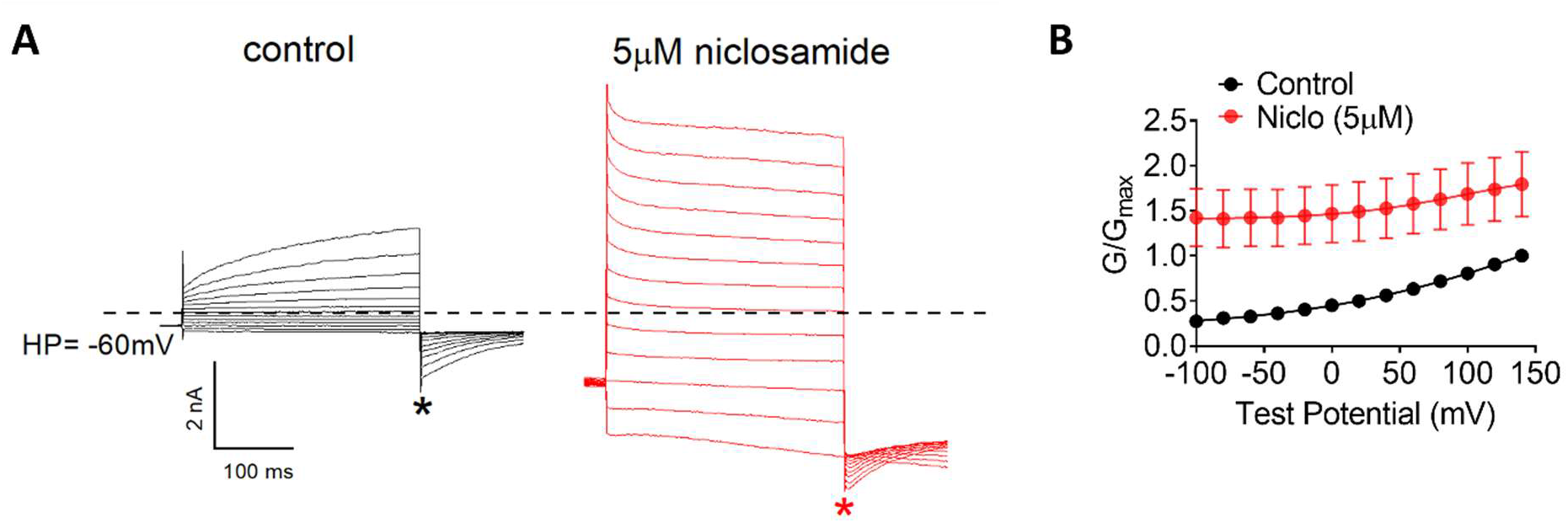
Niclosamide potentiates TMEM16A ‘ac’ splicing isoform overexpressed in HEK293T cells. **A,** Representative whole-cell current traces in the absence (control) or presence of 5 µM niclosamide induced by a voltage-step protocol from −100 mV to +140 mV with 20-mV increments. Holding potential was −60 mV. **B,** *G-V* relationship of TMEM16A currents recorded in A. All tail currents were normalized to the control tail current elicited by a +140 mV depolarization pulse. Error bars represent SEM, n=5.

Because Ca^2+^-activated Cl^-^ efflux via TMEM16A is a potent driving force for water flux and cell volume regulation (Almaça et al., 2009; Takayama et al., 2015). we examined niclosamide’s effect on cell volume. In the absence of niclosamide, no obvious cell volume change occurred during 2 min of whole-cell recording in HEK293T cells overexpresssing TMEM16A at a holding voltage of −60 mV with 500 nM intracellular Ca^2+^ (Fig. 3A and Video S1). This is consistent with little or no opening of TMEM16A under this condition (Fig. 1B-C). On the other hand, 5 µM niclosamide induced dramatic cell shrinkage (Fig. 3B and Video S2). The niclosamide-induced shrinkage is likely due to its potentiation effect on TMEM16A, which promotes Cl^-^ and water efflux. In stark contrast, 5 µM niclosamide had no obvious effect on the volume of untransfected HEK293T cells lacking TMEM16A overexpression (Fig. 3C and Video S3). Our cell volume experiments thus further support that niclosamide potentiates exogenous TMEM16A.

**Figure 3.**
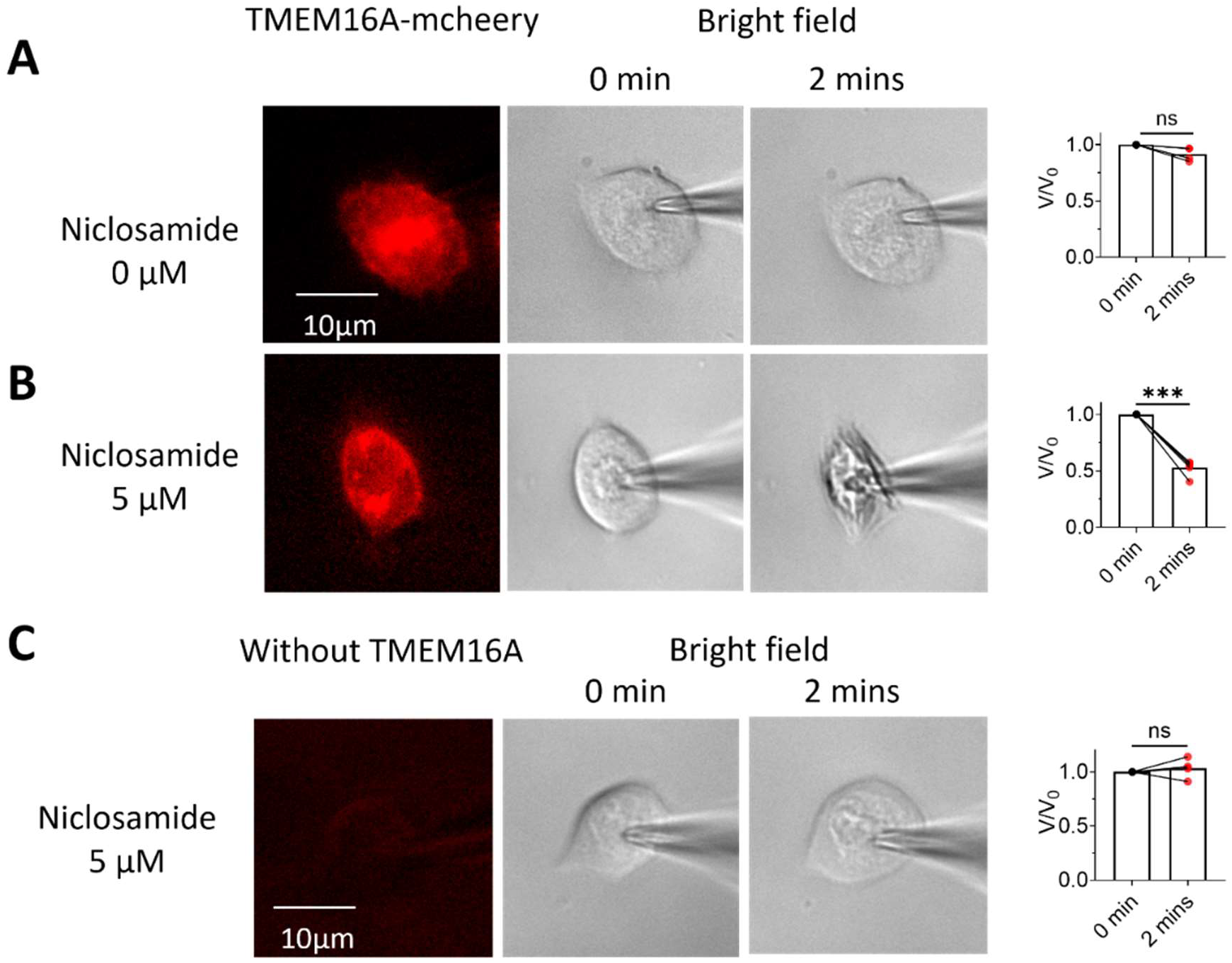
Niclosamide stimulates cell shrinkage in HEK293T cells expressing TMEM16A. **A-B.** Representative images (left) of cell volume changes of voltage-clamped HEK293T cells overexpressing TMEM16A-mCherry in the absence (A) and presence (B) of 5 µM niclosamide. Right: comparison of cell volume changes before and after niclosamide. Statistical analysis was done with Student’s t test (ns means no significance, *** means p<0.001, n=4 for A and n=5 for B). **C**. Representative images (left) of the cell volume changes of the voltage clamped HEK293T cells without TMEM16A overexpression treated with 5 µM niclosamide. Right: comparison of cell volume changes before and after niclosamide. Statistical analysis was done with Student’s t test (ns means no significance, n=4). All experiments were done under whole-cell patch clamp configuration with holding potential at −60 mV and intracellular free Ca^2+^ of 500 nM.

### Niclosamide potentiates TMEM16A from the extracelluar side

Niclosamide has been reported to exhibit multiple off-target effects such as interfering with intracellular Ca^2+^ signaling and inhibiting G protein coupled receptors (GPCRs) (De Filippo et al., 2017; Genovese et al., 2023). To avoid effects of niclosamide on intracellular signaling, we conducted excised patch clamp experiments under outside-out and inside-out configurations. Consistent with our whole-cell patch clamp recording, extracellular application of 5 µM niclosamide to outside-out patches markedly potentiates TMEM16A activation and its effect is reversible (Fig. 4A-B). In contrast, intracellular application of 5 µM niclosamide to inside-out patches has no effect on TMEM16A current (Fig. 4C-D). Our excised patch clamp experiments thus demonstrate that niclosamide directly potentiates TMEM16A from extracellular side and not the intracellular side.

**Figure 4.**
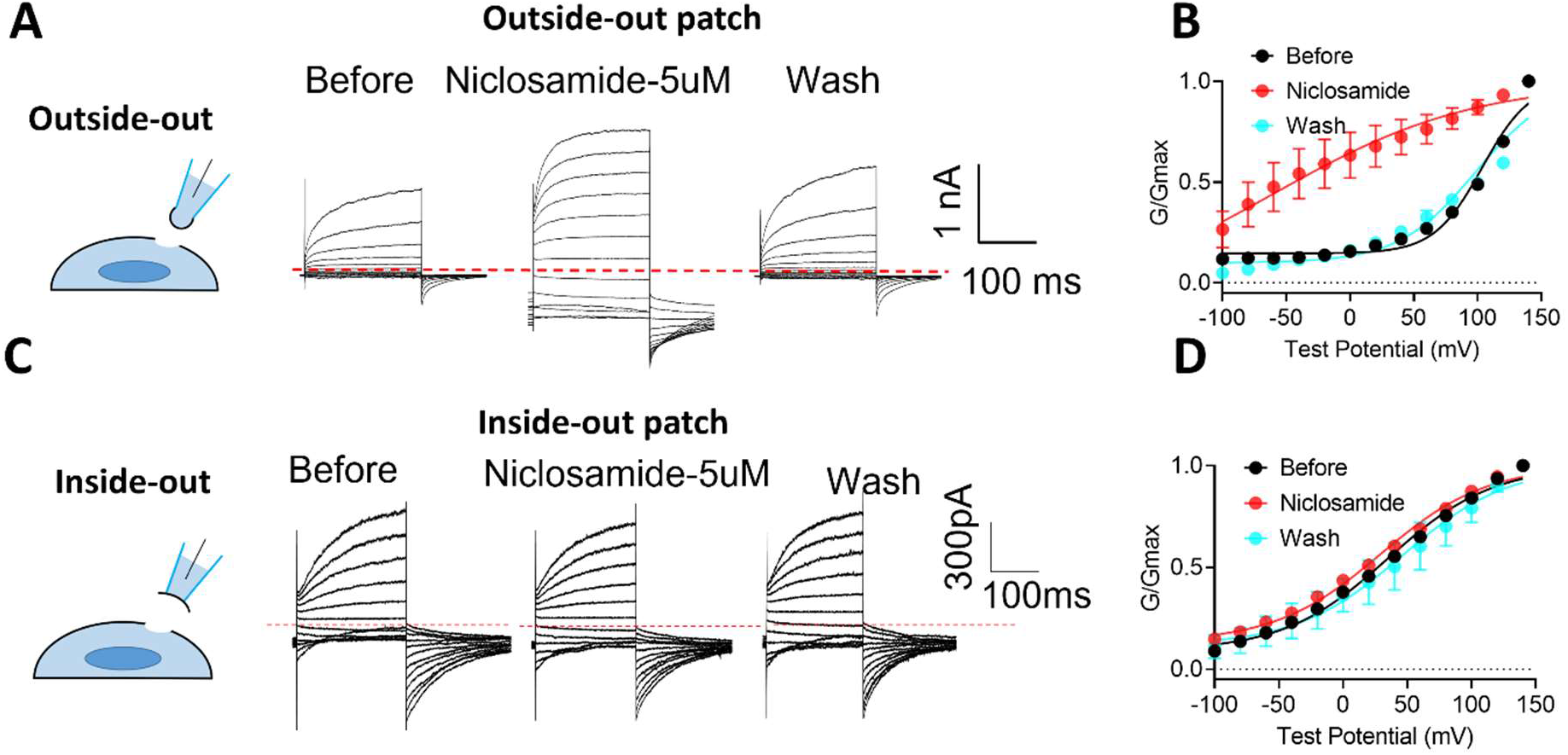
Niclosamide directly targets TMEM16A from extracelluar side. **A,** Representative currents recorded with outside-out patches from HEK293T cells overexpressing TMEM16A. Intracellular solution contained 200 nM free Ca^2+^. The current was elicited by a voltage step protocol from −100 mV to +140 mV with 20-mV increments. Holding potential was −60 mV. The patch clamp configuration is shown on the left. **B,** *G-V* relation of the currents recorded in A. Error bars represent SEM, n=5. **C,** Representative currents recorded in HEK293T cells overexpressing TMEM16A with inside-out patches. The current was elicited by a voltage step protocol from −100 mV to +140 mV with 20-mV increments. Holding potential was −60 mV. The intracellular solution contained 387 nM free Ca^2+^. The patch clamp configuration is shown on the left. **D,** *G-V* relation of the currents recorded in C. The tail currents were normalized to that of the control at +140 mV. Error bars represent SEM, n=4.

### Identification of putative niclosamide binding residues

To dissect the molecular underpinning of niclosamide-mediated TMEM16A potentiation, we sought to identify the niclosamide binding site on the extracellular side of the channel. Through the combination of molecular docking and all-atom molecular dynamics (MD) simulation (see Methods), we identified an extracellular site that favors niclosamide binding (Fig. 5A). The highest scoring pose has a docking score of −8.837. The binding site is a very hydrophobic pocket near the extracellular side of the membrane that is lined with hydrophobic and positively charged amino acids from transmembrane helices (TM) 5, 9, 10. Niclosamide is predicted to have strong interactions with R605 and F601 located at the extracellular end of the TM5. The sidechains of these residues face towards the dimer interface and away from the ion permeation pathway (Fig. 5A). The NH2 group of R605 makes salt bridge contacts with the carboxy oxygen and hydrogen bonds with the carbonyl oxygen of the niclosamide hydroxybenamide moiety. F601 makes a pi-pi stack with the hydroxybenzamide ring and F781 on TM 9 makes a pi-pi stack with the nitrophenyl group.

**Figure 5.**
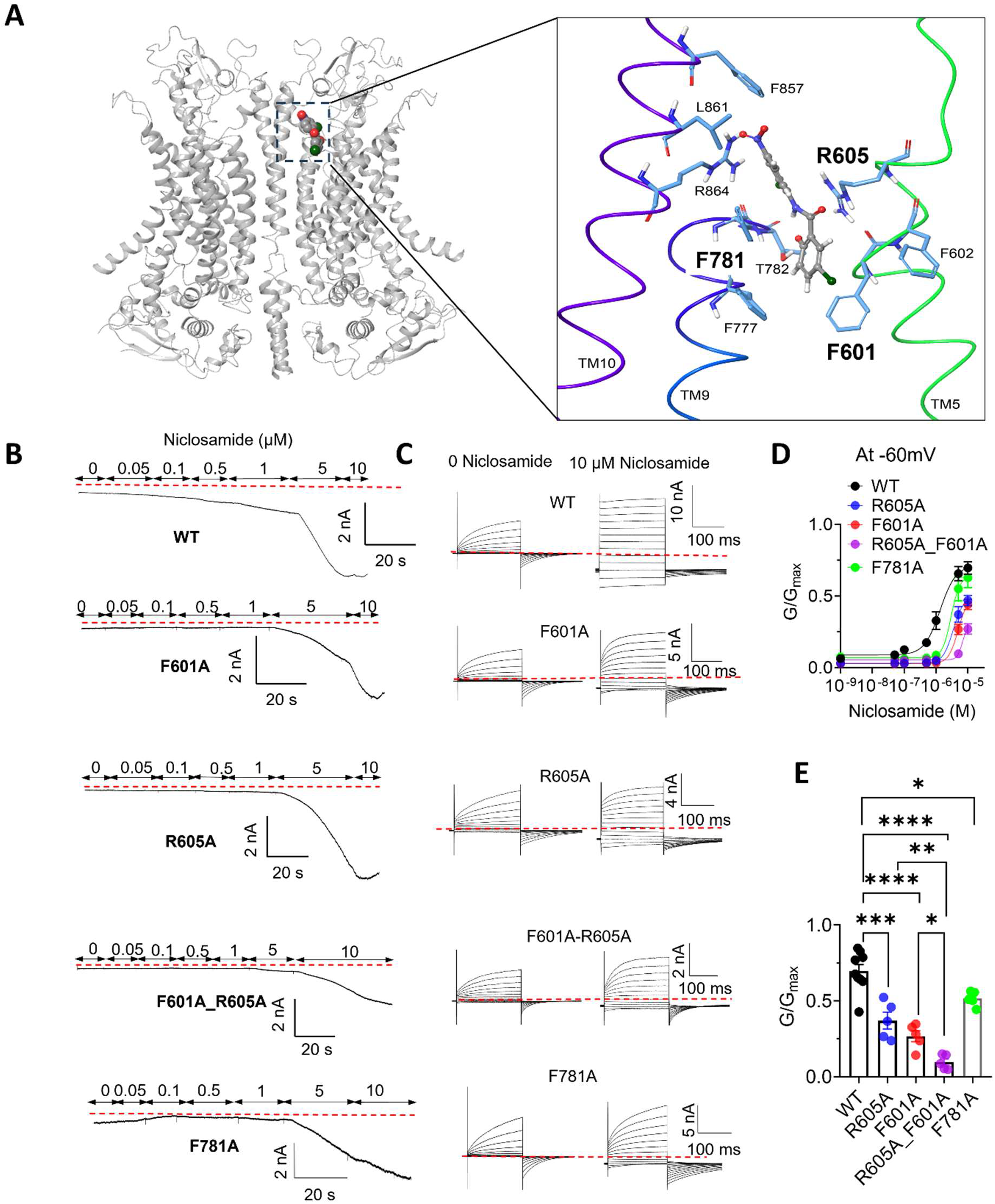
Mutations of a putative niclosamide binding site attenuate the niclosamide potentiation effect. **A.** Computational prediction of a putative niclosamide binding sites in TMEM16A extracellular side (PBD 5OYB). The zoom-in view of potential niclosamide interacting residues is shown on the right. **B.** Representative whole-cell current traces of WT and mutant TMEM16A channels elicited by a gap-free protocol with holding potential of −60 mV in response to different concentrations of niclosamide. Intracellular free Ca^2+^ was 500 nM. **C.** Representative current traces of WT and mutants with and without 10 µM niclosamide. The current from the same patches in (B) were elicited by a voltage-step protocol from −100 mV to +140 mV with 20 mV increments. Holding potential was −60 mV. **D.** Dose-response curve of the niclosamide effect. The currents in (B) were normalized to the maximum tail current of the same patch elicited by a +140 mV test pulse in the presence of 10 µM niclosamide. Data were derived from (C).. **E.** Effect of mutations on 10 µM niclosamide-induced TMEM16A potentiation at −60 mV. Data were derived from (D). Statistical analysis was done by one-way ANOVA. *: p<0.05, **: p<0.01, ***: p<0.001, ****: p<0.0001, n=9 for WT and n=5 for the mutations.

To test the significance of these potential interactions, we substituted alanine for each of these residues and measured the apparent sensitivityof these mutants to niclosamide using whole-cell patch clamp (Fig. 5B-D). For the wildtype (WT) channel, niclosamide dose-dependently potentiates TMEM16A current. The lowest concentration of niclosamide to produce obvious channel activation occurs at ∼0.1 µM. The EC_50_ is 1.54 ± 0.46 µM (Fig. 5B). The F601A, R605A and F781A mutations drastically attenuate niclosamide potentiation. They increase the minimum niclosamide concentration for TMEM16F activation by >10-fold (> 1 µM) and produce a rightward shift the niclosamide dose-respsonse curves at −60 mV with EC_50_’s of 7.19 ± 0.66 µM, 5.66 ± 0.91 µM and 3.07 ± 0.75 µM, respectively (Fig. 5B-C). We further quantified niclosamide-induced potentiation by normalizing the 10 µM niclosamide-elicited current at −60 mV (Fig. 5B) to the TMEM16A tail conductance (G) at −60 mV elicted by 10 µM niclosamide and a depolarizing test pulse to +140 mV (G_max_, Fig. 5C). The WT TMEM16A current in the presence of 10 µM niclosamide is approximately 70% of G_max_, whereas this percentage was decreased significantly to 30-40% for the single mutations F601A and R605A (Fig. 5D-E). The F601A-R605A double mutant channel shows the most rightward-shifted niclosamide dose response curve with an EC_50_ of 79.60 ± 3.67 µM and a potentation of only ∼10% of G_max_ (Fig. 5D-E), suggesting that the effects of mutations on these two residues are additive. The alanine mutations have negibile effects on the voltage-dependent activation of TMEM16A, as evidenced by the identical conductance-voltage relationship (*G-V*) and the similar half-activation voltage (Fig. 6), indicating that the mutational effect on reducing niclosamide potentiation is not due to their influence on channel activation.

**Figure 6.**
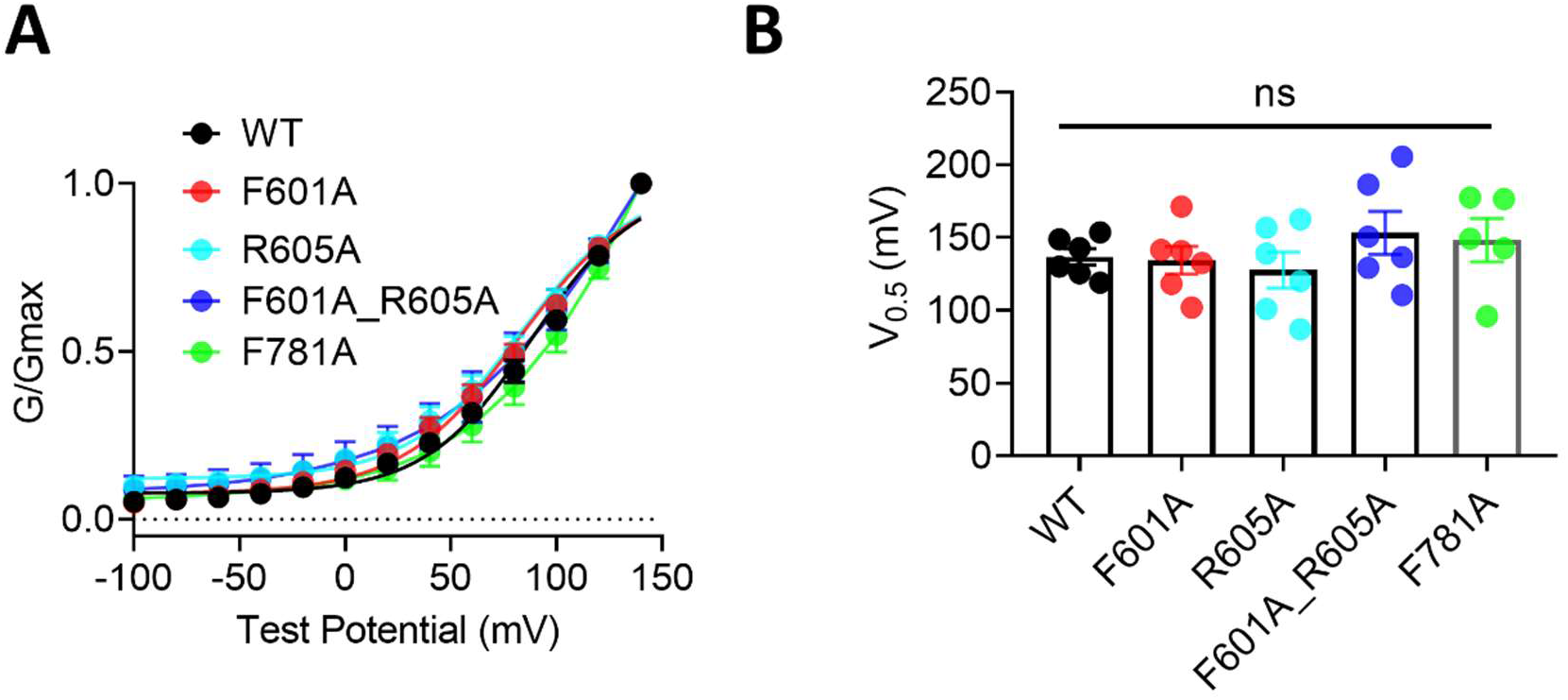
The mutations of the putative niclosamide binding site residues do not alter TMEM16A activation. **A,** *G-V* relationship of TMEM16A wildtype (WT) and mutant channels recorded under whole-cell configuration with 500 nM free intracellular Ca^2+^. Error bars represent SEM, n=6 for each construct. All tail current was normalized to the tail current elicited by a +140 mV voltage pulse. **B,** V_0.5_ of currents derived from the *G-V* relations in B. Statistical analysis was done with one-way ANOVA (ns: not significant, n=5-6).

Potentiation by niclosamide is accompanied by a significant decrease in the outward rectification that is characteristic of TMEM16A current observed with sub-micromolar intracellular Ca^2+^ (Figs. 1B and 5C). The observation that both the single and double mutations largely preserve TMEM16A outward rectifcation (Fig. 5C), further supports the conclusion that the alanine mutations of F601 and R605 attenuate niclosamide-mediated TMEM16A potentiation. Taken together, our computational and functional tests identified a putative niclosamide binding site on the extracellar side of TMEM16A. Residues F601, R605 and F781 within the putative binding site provide potential coordinates for niclosamide binding.

### Niclosamide potentiates endogenous TMEM16A in vascular smooth muscle cells

TMEM16A is abundantly expressed in vascular smooth muscle cells (VSMCs) and plays a crucial role in regulating VSMC membrane potential, activation of the voltage-gated calcium channels (VGCCs) and subsequent VSMC contraction and vascular tone (Huang et al., 2009; Huang et al., 2012; Hawn et al., 2021; Wray et al., 2021). To examine the niclosamide effect on endogenous TMEM16A, we studied VSMCs derived from the aorta of Acta2-GCaMP8.1/mVermilion mice, which express the genetically encoded Ca^2+^ sensor GCaMP8.1 in VSMCs under the control of the *Acta2* locus promotor (Lee et al., 2021). The GCaMP8.1 and mVermilion postitive VSMCs were recorded under whole-cell configuration with 140 mM TEA-Cl to block K^+^ channels. Endogenous CaCC was activated by 500 nM Ca^2+^ in the internal (pipette) solution. Under this condition, we recorded a CaCC current with outward rectification that is a characteristic of TMEM16A. This current was greatly potentiated and its outward rectification was largely diminished by 5 μM extracellular niclosamide (Fig. 7A-D). Niclosamide-mediated potentation of the endogenous TMEM16A current is fully reversable.

**Figure 7.**
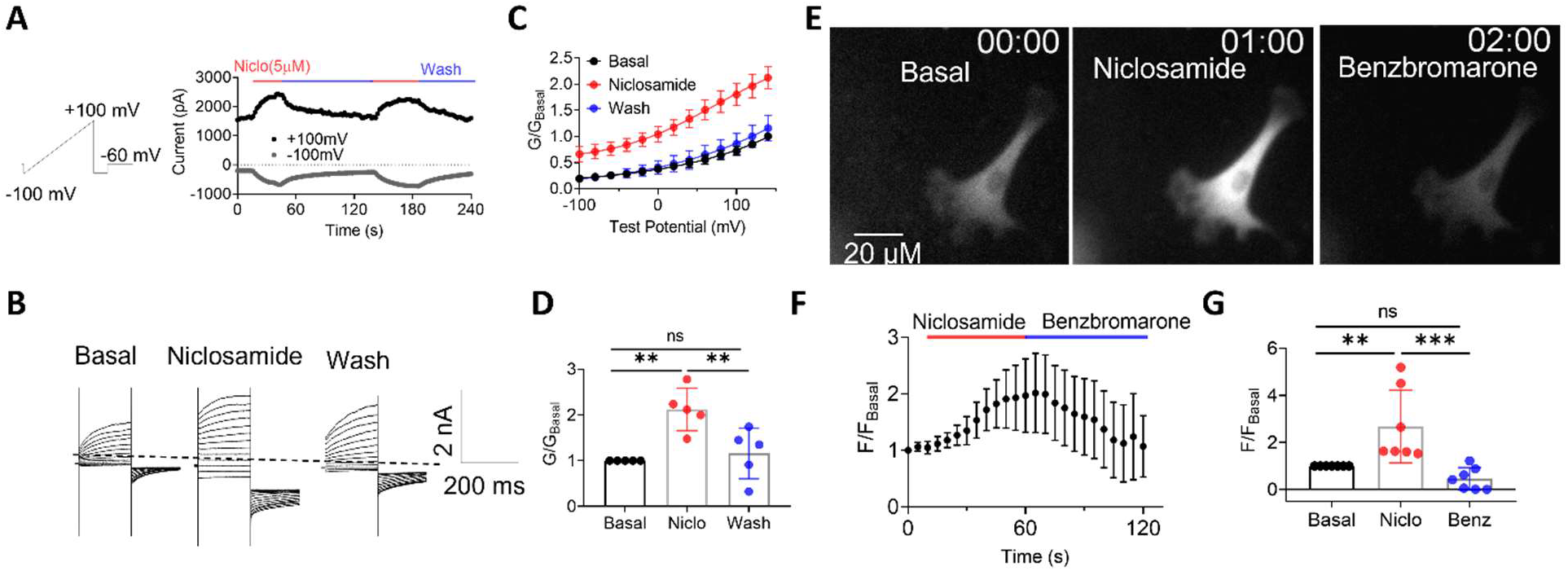
Niclosamide potentiates endogenous CaCC in murine aortic vascular smooth muscle cells (SMCs). **A,** Representative whole-cell CaCC current recorded from aortic SMCs in response to 5 µM niclosamide (Niclo). Intracellular solution contained 500 nM free Ca^2+^. Voltage protocol is shown on the left. **B,** Representative current traces. Voltage steps from-100 mV to +140 mV in 20-mV increments.Holding potential was −60 mV. **C,** *G-V* relationship of the current recorded in B. The tail currents were normalized to the tail current elicited by a +140 mV voltage step before niclosamide (G_Basal_). **D**, Comparison of normalized tail current elicited by a +140 mV voltage step with and without niclosamide. Statistical analysis was done by one-way ANOVA. **: p<0.01, n=5. ns: not siginifcant. **E,** Representative GCaMP8.1 images of intracellular Ca^2+^ changes in response to 5 μM niclosamide and 100 μM benzbromarone, a TMEM16A inhibitor. The aortic SMCs were isolated from the Acta2-GCaMP8.1/mVermilion mice. **F,** Time-course of Ca^2+^ dynamics in response to niclosamide and benzbromarone. Error bars represents SEM from 7 cells. **G,** Normalized Ca^2+^ signal (to basal level) after application of niclosamide and benzbromarone. Statistical analysis was done using one-way ANOVA. **: p<0.01, ***: p<0.001, ns: not significant. n=7.

To further examine the impact of niclosamide-mediated potentiation of endogenous TMEM16A on VSMC function, we conducted Ca^2+^ imaging experiments on the VSMCs from Acta2-GCaMP8.1/mVermilion mice. We found that 5 μM niclosamide robustly increased intracellular Ca^2+^. Because this acute Ca^2+^increase was suppressed by benzbromarone (Fig. 7E-G and Video S4), which is a TMEM16A inhibitor (Huang et al., 2012; Sung et al., 2016), we conclude that niclosamide stimulates TMEM16A, whose Cl^-^ conductance leads to membrane depolarization, VGCC activation, and subsequent increases in intracellular Ca^2+^.

### Niclosamide triggers vasoconstriction

Our finding that niclosamide elevates cytosolic Ca^2+^ in VSMCs suggests that it could induce vasoconstriction. To test this hypothesis, we used *ex vivo* pressure myograph to quantify the effect of niclosamide on contractility of the third order mesenteric arteries from C57BL/6J mice. We first examined the niclosamide effect on equilibrated, non-treated vessels. We find that 5 μM niclosamide induces ∼ 40% vessel contraction as compared to 10 μM phenylephrine (PE) (Fig. 8A-C). In addition, we find that 5 μM niclosamide has a negligible vasodilatory effect on the PE-contracted mesenteric arteries (Fig. 8D-F). In contrast, benzbromarone produces almost complete vasodilation of the PE-contracted artery comparable to the level of relaxation produced by acetylcholine (ACh) (Fig. 8D-F). To further evaluate the effects of niclosamide on vasoconstriction, we applied 5 μM niclosamide to the ACh-treated mesenteric arteries. We find that niclosamide results in vesoconstriction, which can be reversed by the administration of benzbromarone (Fig. 8G-I). Consistent with our observations that niclosamide potentiates exogenous and endogenous TMEM16A CaCC (Fig. 1 and Fig. 7) and triggers Ca^2+^ increases in VSMCs (Fig. 7), our myograph measurements demonstrate that acute niclosamide administration causes vasoconstriction by potentiating VSMC TMEM16A and intracellular calcium increase.

**Figure 8.**
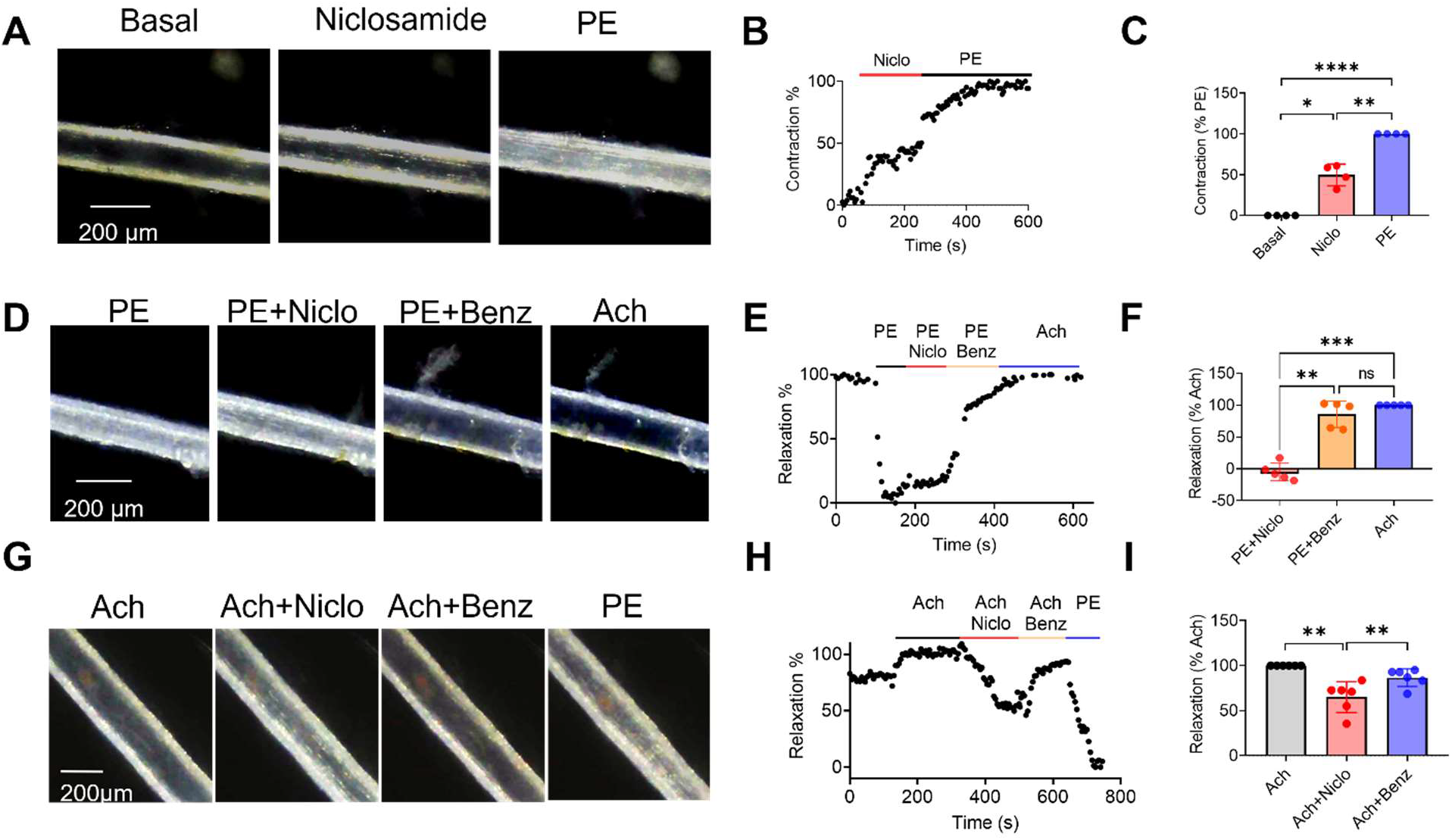
Niclosamide causes vasoconstriction instead of vasodilation. **A,** Representative pressure myograph images of the mesenteric artery in response to sequential application of compounds, including 5 µM niclosamide (Niclo) and 10 µM phenylephrine (PE). **B,** Relative outer diameter changes of the mesenteric artery in A. **C,** Comparison of drug-induced mesenteric artery constriction. Data was calculated as constriction percentages normalized to the PE-induced complete constriction (see Methods for details). Statistical analysis was done by one-way ANOVA. *: p<0.05, **: p<0.01,**** p<0.0001, n=4. **D,** Representative presure myograph images of the PE pre-treated mesenteric artery in response to sequential application of compounds, including 10 µM PE, 10 µM PE + 5 µM Niclo, 10 µM PE + 100 µM benzbromarone (Benz), and 10 µM acetycholine (ACh). **E,** Relative outer diameter changes of the mesenteric artery in D. **F,** Comparison of drug-induced mesenteric artery dialation. Data was calculated as relaxation percentages normalized to the Ach-induced complete vasodilation (see Methods for details). Statistical analysis was done by one-way ANOVA. **: p<0.01, ***: p<0.001, n=5. **G,** Representative presure myograph images of the Ach-treated mesenteric artery in response to sequential application of compounds, including 10 µM ACh, 10 µM ACh + 5 µM Niclo, 10 µM ACh + 100 µM Benz, and 10 µM PE. **H,** Relative outer diameter changes of the mesenteric artery in G. **I,** Comparison of drug-induced mesenteric artery dialation. Data was normalized to the ACh group. Statistical analysis was done using one-way ANOVA. **: p<0.01, n=6.

## DISCUSSION

This study provides evidence at molecular, cellular, and tissue levels that niclosamide is a potent TMEM16A potentiator. Niclosamide potentiates TMEM16A currents in HEK293 cells overexpressing TMEM16A from the extracellular side. In VSMCs expressing endogenous TMEM16A, niclosamide stimulates TMEM16A and depolarizes the cell, which increases cytosolic Ca^2+^ and produces vasoconstriction. Consistent with a recent report (Danahay et al., 2023), we find no indication that niclosamide inhibits TMEM16A CaCC at physiological Ca^2+^ concentration and voltages.

We have examined the effect of niclosamide in both whole-cell and excised patch clamp configurations. First, our whole-cell patch clamp recording under well-buffered physiological calcium concentration shows extracellular niclosamide potentiates TMEM16A current in an acute and reversible fashion (Fig. 1). Second, we observed niclosamide-induced cell shrinkage when TMEM16A-expressing cells were held at −60 mV under whole-cell patch clamp configuration (Fig. 3). This indicates that niclosamide induces cell dehydration driven by chloride efflux through TMEM16A, supporting the conclusion that niclosamide potentiates the channel. Third, our excised patch clamp experiments demonstrate that niclosamide potentiates TMEM16A from the extracellular side instead of intracellular side (Fig. 4). With membrane patches in the outside-out configuration, niclosamide potently and reversibly poentiates TMEM16A. As this configuration eliminates the potential off-target effects of niclosamide on intracellular signaling, our excised patch clamp experiments explicitly show that niclosamide directly potentiates TMEM16A from extracellular side. Fourth, three mutations, F601A, R605A and F781A, in the putative niclosamide binding site on the extracellular side significantly attentuate niclosamide’s potentiation effect, further supporting that niclosamide directly acts on TMEM16A to potentiate the channel. Fifth, our whole-cell patch clamp recording on vascular SMCs shows that extracellular niclosamide also potentiates endogeous TMEM16A CaCC (Fig. 7).

The identification of the putative niclosamide binding site on the extracellular side of TMEM16A supports that niclosamide indeed directly acts on the channel. The binding pocket resides close to the dimer cavity but slightly away from the ion permeation pathway. It is likely that niclosamide binding to this putative site triggers allosteric arrangements of the channel gate, thereby potentiating TMEM16A activation. Future studies are needed to understand the mechanism underlying this drug-induced TMEM16A potentiation. The binding pocket can serve as a potential drug binding site to conduct *in silico* drug screening to identify novel allosteric gating modulators for TMEM16A.

Our data contrast with some prior studies suggesting that niclosamide acts as a TMEM16A antagonist (Miner et al., 2019; Henckels et al., 2020). We find no indication that niclosamide inhibits TMEM16A. A recent study of different TMEM16A isoforms (Danahay et al., 2023) and our independent patch clamp characterizations in two laboratories with two different TMEM16A isoforms rule out the possibility that the reported inhibitory effects of niclosamide is derived from different splicing isoforms.

Previous studies also showed inconsistent niclosamide effects at the tissue level (Li et al., 2017; Miner et al., 2019; Danahay et al., 2020; Danahay et al., 2023). The discrepancy may also derive from niclosamide’s broad molecular targets. Although niclosamide was shown to relax pre-contracted human bronchial rings (Miner 2019 and Danahay 2020), it pardoxically constricts human pulmonary arteries (Danahay 2020), consistent with the vasoconstriction effect in murine mesenteric ateries observed in our study (Fig. 8). It is unclear why the airway and the ateries show opposite responses to niclosamide. It is likely that niclosamide may target different sets of proteins in the two different types of SMCs.

In conclusion, our current study provides compelling evidence to show that niclosamide directly potentiates TMEM16A at the molecular level under physiological conditions. Its potentiation effect on TMEM16A in vascular SMCs may partially contribute to the observed niclosamide effect on promoting vasoconstriction. Overall, niclosamide is a compound with multifaceted effects on various protein targets. When attempting to repurpose niclosamide as a TMEM16A antagonist to treat asthma, COPD, and hypertension, its potentiation effect on TMEM16 CaCCs and its broad impacts on different targets need to be carefully evaluated to avoid triggering severe side effects in human patients.

## METHODS

### Reagents

Niclosamide (Sigma Aldrich, Lot#: 047M4946V), phenylephrine hydrochloride (PE) (Sigma Aldrich, CAT#: P6126), acetylcholine chloride (Ach) (Sigma Aldrich, Lot #:BCCD7863), amphotericin B (Gibco, Lot #: 2419073), and collagenase type 2 (Worthington, CAT#: LS004177). Other reagents were purchased from Sigma Aldrich unless otherwise specified.

### Mouse models

Mouse handling and usage were carried out in strict compliance with the protocol approved by the Institutional Animal Care and Use Committee at Duke University, in accordance with National Institute of Health guidelines. C57BL6/J (stock # 000664) mice and Acta2-GCaMP8.1/mVermilion (stock #032887) (Lee et al., 2021) mice expressing calcium sensor GCaMP8.1 and mVermilion reporter in SMCs were purchased from the Jackson Laboratory.

### Pressure myograph

The experiments followed the procedures as previously described (Shahid and Buys, 2013; Rode et al., 2017; Turner et al., 2019). Briefly, the third order mesenteric arteries were dissected and placed immediately into ice cold HEPES-PSS solution (125 mM NaCl, 3.8 mM KCl, 1.2 mM CaCl_2_, 25 mM NaHCO_3_, 1.2 mM KH_2_PO_4_, 1.5 mM MgSO_4_, 0.02 mM EDTA and 8 mM D-glucose, pH 7.4) pre-equilibrated with 95% O_2_/5% CO_2_ for 15 min. Small segments of 3–5 mm length with no branching were cut from the mesenteric artery and mounted on glass cannulas in a pressure myograph (Model 110p, Danish Myo Technology A/S, Denmark). Two pressure transducers (P1 on the right and P2 on the left side) are built in to monitor the pressure within the artery lumen. The vessels were equilibrated at 60 mmHg and 37 °C for at least 45 min. The bath solution was changed once with pre-warmed HEPES during equilibration. The viability of the vessel and integrity of the endothelial cells was assessed by addition of 10 µM phenylephrine (PE) to constrict and 10 µM acetylcholine (ACh) to relax to vessel. Only the arteries that were constricted by PE and dilated by ACh were included. The vessel pressure was maintained at constant 60 mmHg throughout the experiments. The outer arterial diameter was monitored using a Cainda WiFi Digital Microscope and analyzed with ImageJ. To change solutions, the myograph bath solution was completely vacuum aspirated and replaced with a new solution from the top. Relaxation percentage was calculated by the formula % relaxation = (D-D_PE_) / (D_Ach_-D_PE_) * 100% and contraction percentage was calculated by formula % contraction = (D_basal_-D) / (D_basal_-D_PE_) * 100%; where D is diameter of the vessel, D_PE_ is the full constricted diameter of the vessel in response to 10 µM PE, D_Ach_ is the fully dilated diameter of the vessel in response to 10 µM Ach and D_basal_ is the diameter of vessel at basal level.

### Isolation and culture of primary aortic smooth muscle cells

Aortic smooth cells were isolated from the Acta2-GCaMP8.1/mVermilion mouse line. GCaMP8.1/mVermilion mice express the GCaMP8.1/mVermilion fusion protein in smooth muscle cells, which allows us to monitor the Ca^2+^ dynamics in response to different regents. Isolation of murine aortic smooth cells was performed as described in detail previously (Adhikari et al., 2015). Briefly, after euthanizing the mouse, aorta was rapidly isolated and transferred to a 6-well plate with complete DMEM solution (10% FBS, 1% penicillin/streptomycin) and Amphotericin B (1µl/ml). The remaining blood was removed by perfusing the aorta using a tuberculin syringe filled with sterile PBS. The surrounding fat tissues were cleaned with fine-tipped forceps. The dissected aorta was then cut into 1-2 mm pieces using fine scissors and placed into 5-mL tubes containing 0.1 mL of complete DMEM with collagenase type 2 (1.42 mg/mL). The aorta was digested at 37 °C (with 5 % CO_2_) for 6 h. After 6h incubation, 3 mL of DMEM plus 10% FBS was added and gently mixed. Cells were centrifuged at 300×*g* for 5 min at room temperature, the medium aspirated, and the pellet washed with 3 mL of DMEM + 10% FBS twice. Cells were incubated at 37 °C with 5 % carbon dioxide, undisturbed for 5 days until full confluence. The cells were sub-cultured by trypsinization (0.25%) for 3-4 minutes and seeded on PLL-coated cover glasses for Ca^2+^ imaging or electrophysiological experiments.

### Ca^2+^ imaging of cultured SMCs

Cultured SMCs were re-seeded on PLL-treated cover glasses. After 24-48 hours, the cells were placed into HEPES-PSS solution (in mM, 125 NaCl, 3.8 KCl, 1.2 CaCl_2_, 25 NaHCO_3_, 1.2 KH_2_PO_4_, 1.5 MgSO_4_, 0.02 EDTA and 8 D-glucose, pH 7.4). The Ca^2+^ signal was monitored using the GFP channel on an Olympus IX73 inverted epifluorescence microscope. Time-lapse images were acquired by a Prime 95B Scientific CMOS Camera (Photometrics) controlled by Metafluor software (Molecular Devices).

### Cell volume change

Untransfected or TMEM16A-transfected HEK-293T cells were patch-clamped under whole-cell configuration and held at −60 mV with 500 nM free Ca^2+^ in the pipet solution. Images were taken every 2 seconds before and after application of 5 µM niclosamide with an Olympus IX73 inverted microscope. Cell volume was quantified by cell area measurement function in ImageJ.

### Cell lines and culture

The generation and validation of the TMEM16A stable cell line and TMEM16F-deficient (knockout [KO]) HEK293T cell line have been reported in our recent studies (Huang et al., 2012; Le et al., 2019; Zhang et al., 2020; Liang and Yang, 2021; Zhang et al., 2022). All the mutants were transiently expressed in TMEM16F-KO HEK293T cells to avoid interference from endogenous TMEM16F. HEK293T cells were cultured with Dulbecco’s modified Eagle’s medium (DMEM; 11995-065; Gibco BRL) supplemented with 10% FBS (F2442; Sigma-Aldrich) and 1% penicillin–streptomycin (15-140-122; Gibco BRL). All cells were cultured in a humidified atmosphere with 5% CO2 at 37°C.

### Mutagenesis and transfection

The murine TMEM16A coding sequence corresponding to the ‘a’ splicing isoform (cDNA 30547439; Open Biosystems) was subcloned in the pEGFP-N1 vector, resulting in eGFP tags on the C terminus. Single point mutations of TMEM16A were generated using QuikChange site-directed mutagenesis and the plasmids were subsequently sequenced. The plasmids were transiently transfected to TMEM16F-KO HEK293T cells using X-tremeGENE9 transfection reagent (Sigma-Aldrich). Cells grew on coverslips coated with poly-L-lysine (Sigma-Aldrich). Medium was changed 6 h after transfection with regular medium (11995-065; Gibco BRL).

Experiments proceeded 24–48 h after transfection. Experiments performed in the Hartzell lab used the TMEM16A (ac) isoform as previously described (Xiao et al. 2011).

Primers for QuikChange mutagenesis are listed in Table 1.

**Table 1.**
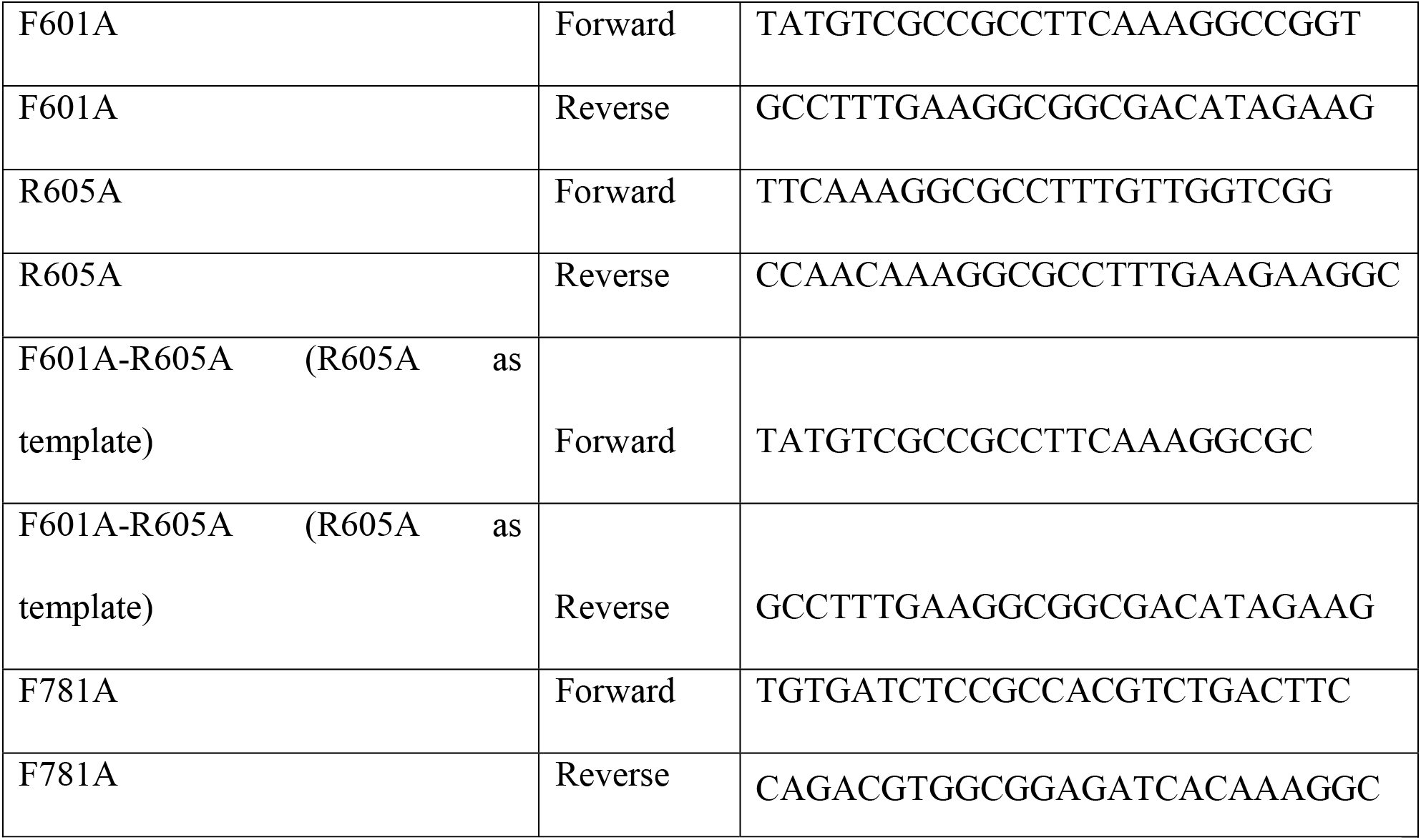
Primers for murine mTMEM16A mutagenesis.

### MD simulation of niclosamide binding

To gain insights into the niclosamide binding site on TMEM16A, we performed docking experiments using the Schrodinger Suite (Schrödinger Release 2022-2: Schrödinger LLC, New York, NY, 2021.) Niclosamide (*5-chloro-N-(2-chloro-4-nitrophenyl)-2-hydroxybenzamide*, PUBCHEM CID 4477) was prepared using Schrodinger LigPrep with the OPLS4 force field (Harder et al., 2016). Ligand tautomeric and ionization and states were generated at pH 7 ± 1 using Epik (Shelley et al., 2007; Greenwood et al., 2010). TMEM16A (PDB: 5OYB) was prepared for docking using the Schrodinger Protein Preparation Wizard in a pH7.4 environment (Sastry et al., 2013). N- and C-termini were capped with N-acetyl and N-methyl amide and missing side chains were added use Prime (Jacobson et al., 2002; Jacobson et al., 2004). H-bond assignments were optimized using PROPKA at pH 7.4 followed by a restrained molecular dynamics minimalization (Olsson et al., 2011; Sondergaard et al., 2011). The niclosamide ligand was docked onto the TMEM16A receptor using Glide. The ligand was docked within a 47 nm^3^ receptor grid that was centered on the transmembrane region of one protomer of the TMEM16A dimer (Friesner et al., 2004; Halgren et al., 2004; Friesner et al., 2006). Induced fit docking was performed using the Induced Fit Protocol (Farid et al., 2006; Sherman et al., 2006a; Sherman et al., 2006b).

### Electrophysiology

All currents were recorded in either inside-out, outside-out or whole-cell configurations 24–48 h after transfection using an Axopatch 200B amplifier (Molecular Devices) and the pClamp software package (Molecular Devices). Glass pipettes were pulled from borosilicate capillaries (Sutter Instruments) and fire-polished using a microforge (Narishge) to reach a resistance of 2–3 MΩ.

For inside-out patch, the pipette solution (external) contained (in mM) 140 NaCl, 10 HEPES, and 1 MgCl_2_, adjusted to pH 7.3 (NaOH), and the bath solution contained 140 NaCl, 10 HEPES, and 5 EGTA, adjusted to pH 7.3 (NaOH). Intracellular (perfusion) solutions with 0.387 μM free Ca^2+^ were made by adding CaCl_2_ to a solution containing 140 mM NaCl, 10 mM HEPES, 5 mM EGTA, and the amount of CaCl_2_ added was calculated using WEBMAXC (https://somapp.ucdmc.ucdavis.edu/pharmacology/bers/maxchelator/) to achieve desired free Ca^2+^.

For outside-out patch, the pipette solution (internal) contained (in mM) 140 mM NaCl, 10 mM HEPES, 5 mM EGTA, and the amount of CaCl_2_ added was calculated using WEBMAXC (https://somapp.ucdmc.ucdavis.edu/pharmacology/bers/maxchelator/) to achieve 200 nM free Ca^2+^. The bath solution contained 140 NaCl, 10 HEPES, and 5 EGTA, adjusted to pH 7.3 (NaOH).

For whole-cell recordings on HEK293T cells, the pipette solution (internal) contained (in mM) 140 CsCl, 1 MgCl_2_, and 10 HEPES, plus CaCl_2_ to obtain the desired free Ca^2+^ concentration using WEBMAXC. For whole-cell recordings on the primary VSMCs cells from Acta2-GCaMP8.1/mVermilion mice, the pipette solution (internal) contained (in mM) 140 TEA-Cl, 1 MgCl_2_, and 10 HEPES, plus CaCl_2_ to obtain the desired free Ca^2+^ concentration using WEBMAXC. Bath solution contained 140 NaCl, 10 HEPES, and 5 EGTA, adjusted to pH 7.3 (NaOH).

Procedures for solution application with/without niclosamide were as used previously (Liang and Yang, 2021). Briefly, a perfusion manifold with 100–200-µm tip was packed with eight PE10 tubes. Each tube was under separate valve control (ALA-VM8; ALA Scientific Instruments), and solution was applied from only one PE10 tube at a time onto the excised patches or whole-cell clamped cells. All experiments were at room temperature (22–25°C). All the chemicals for solution preparation were obtained from Sigma-Aldrich including niclosamide.

### Data Analysis

*G-V* curves were constructed from tail currents measured 200-400 μs after repolarization. For the *G-V* curves obtained from the same patch, the conductance was normalized to the tail current at +140 mV before niclosamide application. Individual *G-V* curves were fitted with a Boltzmann function,

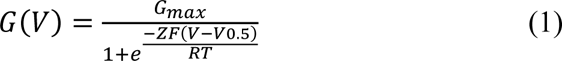

where *G_max_* denotes the fitted value for maximal conductance, *V_0.5_* denotes the voltage of half maximal activation of conductance, *z* denotes the net charge moved across the membrane during the transition from the closed to the open state and *F* denotes the faraday constant.

For binding site mutation analysis, the currents were generated by a gap-free protocol holding at −60 mV in response to desired niclosamide concentrations. Right after the gap-free protocol, a voltage-step protocol was applied from −100 mV to +140 mV in the presence of highest niclosamide concentration (10 µM). All the currents at −60 mV in response to different niclosamide concentrations were then normalized to the maximum tail current at +140 mV in the presence of 10 µM niclosamide (G_Max_).

Dose response curves were fitted to the Hill equation,

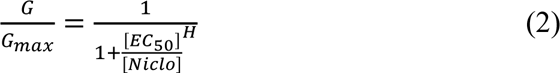

where *G/G_max_* denotes current normalized to the max current elicited by 10 µM niclosamide. [Niclo] denotes niclosamide concentration, *H* denotes Hill coefficient, and *EC_50_* denotes the half-maximal activation concentration of Ca^2+^.

## ACKNOWLEDGEMENT

This work was supported by the NIH-DP2GM126898 grant (awarded to H.Y.), NIH R01-132598 to HCH) and AHA postdoctoral fellowship to P.L. (#903807).

## AUTHOR CONTRIBUTIONS

H.Y. and H.C.H. conceived and supervised the project. H.Y., H.C.H. and P.L. designed the research. P.L. and K.Y. performed electrophysiology experiments; Y.S.W. did mutagenesis; H.C.H. performed computation. P.L., K.Y. and H.C.H analyzed the data. P.L., H.C.H. and H.Y. wrote the manuscript with input from all authors.

